# A high-content *in vivo* screen to identify microRNA epistasis in the repopulating mouse liver

**DOI:** 10.1101/664847

**Authors:** Adam M. Zahm, Amber W. Wang, Yue J. Wang, Jonathan Schug, Kirk J. Wangensteen, Klaus H. Kaestner

**Author notes:** All work performed in the Department of Genetics, University of Pennsylvania, Philadelphia, Pennsylvania. **Abbreviations:** AGO2 (Argonaute 2), FAH (fumarylacetoacetate hydrolase), FLuc (firefly luciferase), FOX (forkhead box), HTI (hereditary tyrosinemia type I), MBS (microRNA-binding site), miRNA (microRNA), NES (Normalized Enrichment Score), NFY (nuclear transcription factor Y), RISC (RNA-induced silencing complex), RLuc (Renilla luciferase), TuD (tough decoy), YBX1 (Y box protein 1). **Correspondence:** Klaus H. Kaestner, 12-126 Smilow Center for Translational Research, 3400 Civic Center Blvd., Philadelphia, PA 19104-6145, Kirk J. Wangensteen, 421 Curie BLVD, BRB 910, Philadelphia, PA 19104. **Disclosures:** The authors have no potential conflicts to disclose. **Transcript Profiling:** Pending. **Author Contributions:** Conceptualization, A.M.Z., K.J.W., and K.H.K.; Methodology, A.M.Z., J.S., and K.J.W.; Formal Analysis, A.M.Z. and Y.J.W.; Investigation, A.M.Z., A.W.W., and K.J.W.; Resources, K.H.K.; Writing - Original Draft, A.M.Z.; Writing - Review & Editing, A.M.Z., A.W.W., Y.J.W., J.S., K.J.W., and K.H.K.; Visualization, A.M.Z. and A.W.W.; Supervision, K.H.K.; Funding Acquisition, K.H.K.

## Abstract

Liver regeneration is impaired in mice with hepatocyte-specific deficiencies in microRNA (miRNA) processing; yet the roles of individual miRNAs or their combinatorial effects in this process are largely unknown. In this study, we sought to identify miRNAs that regulate hepatocyte repopulation following toxic liver injury in a high-throughput manner using the *Fah*^*−/−*^ mouse. We constructed plasmid pools encoding over 30,000 tough decoy (TuD) miRNA inhibitors designed to target hepatocyte miRNAs in a pairwise manner. Plasmid libraries were delivered to hepatocytes of *Fah*^*−/−*^ mice at the time of liver injury via hydrodynamic tail vein injection and integrated transgene-containing transposons were quantified following repopulation via high-throughput sequencing. Changes in polysome-bound transcripts following miRNA inhibition were determined using translating ribosome affinity purification followed by high-throughput sequencing. Analysis of TuD abundance in hepatocyte genomic DNA and input plasmid pools identified several thousand miRNA inhibitors that were significantly altered following repopulation. We classified a subset of miRNA-binding sites (MBSs) as having strong effect on liver repopulation, thus implicating the targeted hepatocyte miRNAs as regulators of this process. Furthermore, we generated a high-content map of pairwise interactions between 171 MBSs and identified both synergistic and redundant effects. Our study highlights the power of higher-order screens to uncover miRNA functions that would go undetected by individual miRNA perturbations, and provides a new paradigm for the study of epistasis of miRNA activities.

## INTRODUCTION

Hepatocytes function at the nexus of numerous essential metabolic pathways and are frequently exposed to environmental toxins and viruses. The ability of the liver parenchyma to restore organ mass in response to tissue injury is highly conserved among vertebrates. In response to injury, local and systemic cytokines and growth factors, such as interleukin 6 and hepatocyte growth factor, signal quiescent hepatocytes to temporarily pause homeostatic functions and re-enter the cell cycle. ^1^ Harnessing this natural ability for regeneration is a topic of clinical importance, as organ transplantation is the only current remedy for fulminant liver failure.

The repopulating hepatocyte transcriptome is highly dynamic^2,3^ but the importance of post-transcriptional gene regulation is poorly understood. MicroRNAs (miRNAs) are a major post-transcriptional regulatory component, controlling mRNA stability and translational efficiency by recruiting the RNA-induced silencing complex (RISC) to target transcripts. ^4^ Previous studies in mice utilizing hepatocyte-specific deletion of Dicer, an obligatory enzyme for miRNA maturation, have shown that miRNAs are required for normal liver development and homeostasis. ^5–7^ Furthermore, mice with a hepatocyte-specific deficiency in miRNA processing exhibit impaired liver regeneration following partial hepatectomy. ^8^ Although a limited number of individual miRNAs have been identified as positive or negative regulators of liver regeneration, no large-scale or combinatorial functional mapping during liver regeneration has been performed to date. Furthermore, interactions between miRNAs during this process are largely unknown.

The direct functional inhibition of endogenous miRNAs is the preferred approach for elucidating their function, as overexpression and miRNA mimetic strategies risk the induction of off-target effects due to supraphysiological concentrations. Tough decoys (‘TuDs’) are a class of potent miRNA inhibitors that can target individual or multiple miRNAs in mammalian cells with high specificity. ^9–11^ TuDs are single-stranded RNAs that contain one or more miRNA-binding sites (MBSs), each designed to base-pair with a mature miRNA, thereby inhibiting interactions of the miRNA with its endogenous mRNA targets (Figure 1A). Evidence that TuDs induce nucleotide trimming and tailing of targeted miRNAs suggests that a single MBS can inactivate multiple miRNA molecules in succession.^12^

**Figure 1.**
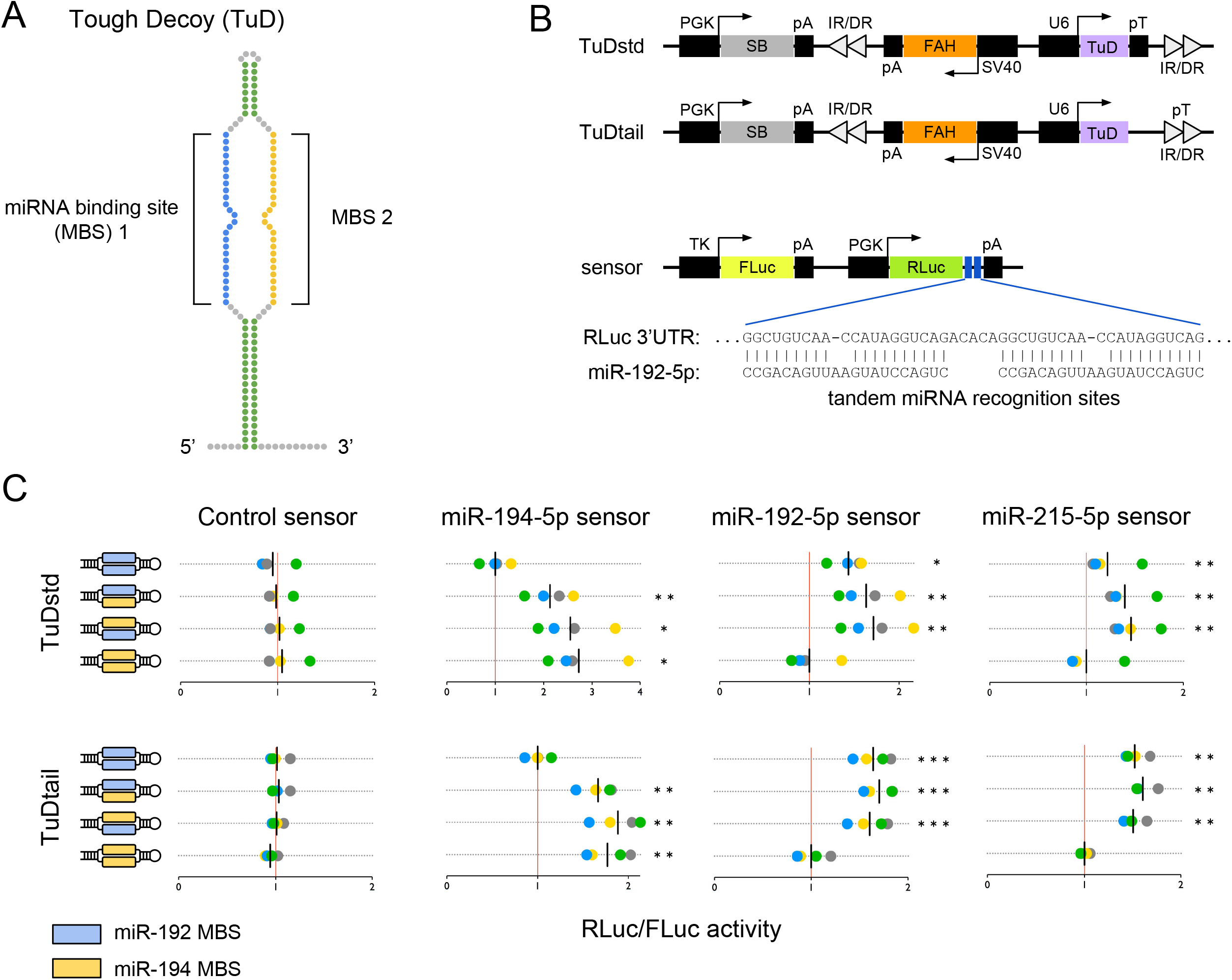
Inhibition of microRNA activity via tough decoy expression. (A) A ssRNA tough decoy (TuD) containing two microRNA-binding sites (MBSs). (B) *Fah* plasmids express TuDs from a U6 promoter as standard hairpins (TuDstd) or with extended 3’ sequences (TuDtail). Plasmids encode sleeping beauty (SB) transposase and inverted repeat/direct repeat (IR/DR) sequences for genomic integration. (C) Dual luciferase assays in mouse Hepa 1-6 cells co-transfected with *Fah* plasmids encoding TuDs directed against the miRNA-194/192/215 family and miRNA sensor plasmids encoding a *Renilla* luciferase (RLuc) cDNA with binding sites for the indicated miRNA in the 3’UTR and a non-targeted firefly luciferase (FLuc) cDNA. The RLuc/FLuc ratios for miRNA sensors are scaled such that the mean of the sensor-specific control TuD is one (red vertical line). Assays were performed in quadruplicate; data points colored by replicate. Black vertical lines indicate the mean luciferase ratio of each TuD. Statistics significance by repeated measures ANOVA with Dunnet’s test for multiple comparisons. *p<0.05; **p<0.01; ***p<0.001.

Partial hepatectomy is a well-characterized and broadly applied model of liver repopulation in rodents but, although a useful tool to study cell cycle entry and progression of mammalian hepatocytes, this approach does not recapitulate the liver injury commonly seen in patients. A pertinent system for studying toxic liver injury and regenerative response is the FAH-deficient (*Fah*^*−/−*^) mouse model of hereditary tyrosinemia type I (HTI), which recapitulates salient features of the condition. ^13^ The *Fah* gene encodes fumarylacetoacetate hydrolase (FAH), which catalyzes the removal of a toxic intermediate of tyrosine catabolism. Individuals with FAH deficiency develop HTI, a condition characterized by toxic injury to the liver and a need for lifelong treatment with the drug nitisinone to inhibit an upstream enzyme to prevent accumulation of the toxic intermediate. A strong selection pressure for cells expressing a *Fah* transgene to survive and expand upon the induction of injury allows co-expressed genes to be screened in parallel for effects on hepatocyte repopulation *in vivo*. ^14–17^ Thus, the *Fah*^*−/−*^ mouse serves as a powerful model of liver repopulation following toxic liver injury.

Utilizing the *Fah*^*−/−*^ model, we developed a large-scale screening platform to map the regulation of liver repopulation by miRNAs following toxic liver injury. We constructed TuD miRNA inhibitors targeting the 171 most abundant RISC-bound hepatocyte miRNAs. By sequential ligation of pooled miRNA inhibitor sequences to a stem-loop scaffold, we created libraries of over 30,000 unique single- and dual-targeting TuDs. We successfully performed the first high-throughput combinatorial inhibition screen to map miRNA function *in vivo*. This approach uncovered previously unknown miRNA regulatory networks active during liver regeneration. The data herein broadens our understanding of hepatocyte post-transcriptional control in the context of liver repopulation and may ultimately aid advancements in therapeutic stimulation of native liver recovery.

## METHODS

### *Fah*^*−/−*^ mouse model of hereditary tyrosinemia type I

C57BL/6j *Fah*^*−/−*^ mice were a gift from Markus Grompe and maintained on 7.5 μg/mL nitisinone in H_2_O until hydrodynamic tail vein injection of plasmids^13^. Animal experiments were approved by the University of Pennsylvania Institute Animal Care and Use Committee (protocol 805623).

### MicroRNA-binding site (MBS) design and TuD library generation

miRNAs were selected using the list of AGO2-bound miRNAs obtained by high-throughput sequencing of RNA isolated by crosslinking immunoprecipitation in repopulating mouse liver following partial hepatectomy. ^3^ One MBS was generated for each confidently-annotated miRNA represented by at least 0.01% of the total reads at any time point during liver regeneration and devoid of a stretch of uracils of length four or more (n=171). Three scrambled-sequence non-targeting MBSs and three MBSs targeting miRNAs detected at less than two reads per million in the Ago2-bound fraction of the quiescent or regenerating mouse liver (miR-1a-1-5p, miR-670-3p, and miR-880-5p) were included as negative controls. MBS oligonucleotides were purchased as ssDNA, annealed and combined to generate a MBS pool containing 177 unique dsDNA MBSs and ligated into pBT264-MBSacceptor. MBS sequences are listed in Supplementary Table 1. After verification, TuDs were ligated to pKT2-Fah-U6-TuDacceptor-pT or pKT2-Fah-U6-TuDacceptor-tail to create the pKT2-Fah-TuDstd (TuDstd) and pKT2-Fah-TuDtail (TuDtail) libraries, respectively. Endotoxin-free TuD library plasmid preparations were obtained using the EndoFree Plasmid Maxi Kit (Qiagen). Individual pKT2-Fah-TuDstd, pKT2-Fah-TuDtail, and pKT2-Fah-eGFP-L10a-TuDstd plasmids containing single- or dual-targeting TuDs were derived by sequentially ligating single MBS inserts into the TuD scaffold of pBT264-MBSaccepto, followed by transfer into pKT2-Fah plasmids.

### Cell culture and dual luciferase assays

Hepa 1-6 mouse hepatoma cells (ATCC CRL-1830) were maintained on culture-treated plastic in Dulbecco’s Modified Eagle Medium supplemented with 10% fetal bovine serum and 100 U/mL penicillin and streptomycin. For luciferase experiments, cells were plated on 24-well tissue culture dishes. Following overnight incubation, cells were co-transfected with 10 ng of pMiRCheck2 miRNA sensor plasmid and 500 ng of individual pKT2-Fah-TuDstd or pKT2-Fah-TuDtail plasmids using Lipofectamine 3000 Reagent (Invitrogen). Cultures were harvested 24 h post-transfection and processed for firefly and *Renilla* luciferase activity using the Dual-Luciferase Reporter Assay System (Promega).

### MicroRNA inhibition in the mouse liver

The TuDstd library or TuDtail library (10 µg in Ringer’s lactated solution; fluid volume was equal to 10% of recipient mouse weight) was delivered to male *Fah*^*−/−*^ mice (age 8-12 weeks) via hydrodynamic tail vein injection. Nitisinone was withdrawn to induce hepatocyte toxicity and repopulation. Four weeks post-injection, mice were euthanized and livers excised. Genomic DNA was isolated from 400 mg of tissue from multiple lobes of each liver and remaining tissue processed for histological analyses. For the TRAP-Seq experiment, pKT2-Fah-eGFP-L10a-TuDstd plasmids expressing either a TuD with two control MBSs (miR-880-5p and scramble-3) or a TuD with a miR-374b-5p MBS in the 3’ position (miR-880-5p and miR-374b-5p) were delivered to male *Fah*^*−/−*^ mice (age 4 months) as above and repopulation induced by. Nitisinone withdrawal. Two weeks post-injection, mice were euthanized and livers isolated for analysis.

### TRAP-Seq

Regenerating hepatocyte-specific messenger RNA was isolated by translating ribosome affinity purification (TRAP) as previously described. ^18^ Briefly, 500 mg of liver homogenate was incubated with magnetic-bound anti-GFP antibodies (Htz-GFP-19F7 and Htz-GFP-19C8, Memorial Sloan-Kettering Monoclonal Antibody Facility, New York, NY). Translating mRNA bound to the GFP-tagged ribosomal subunit protein L10a was then pulled-down and purified (Absolutely RNA Microprep Kit, Agilent Technologies, Wilmington, DE). RNA-Seq libraries were prepared from 1 μg of purified RNA using the NEBNext Ultra RNA Library Prep Kit for Illumina with NEBNext Poly(A) mRNA Magnetic Isolation Module (New England BioLabs). Sequencing was performed on an Illumina NextSeq 500 (75 cycles). Following alignment to the GRCm38/mm10 genome assembly using STAR v2.5.2a, raw read counts were converted to transcripts per kilobase million (TPM).

### Immunohistology

Paraffin sections were labeled with antibodies according to standard procedures. For the detection of FAH, sections were incubated overnight with a rabbit anti-FAH antibody diluted 1:200 in PBS with 10% FBS. GFP was detected by overnight incubation with a goat anti-GFP antibody diluted 1:500.

### Differential TuD abundance

After de-multiplexing of high-throughput sequencing reads, TuD sequence read counts were generated as follows. We generated a custom TuD dictionary from a dictionary of 31,329 potential TuD sequences by removing stem sequence, trimming to 75 nucleotides, and removing invariable center loop sequence. The dictionary was expanded to allow a single mismatch across the trimmed sequences. Reads corresponding to MBS1 and MBS2 were concatenated and mapped to the TuD name (MSB pair). Using our strict mismatch criteria, approximately 95% of reads mapped to a specific TuD sequence.

We used the mapped raw TuD sequencing reads to determine differential TuD sequence abundance following liver repopulation using the The DESeq2 algorithm^19^. A Benjamini-Hochberg adjusted p-value (FDR) less than 0.05 was considered statistically significant.

## RESULTS

### Inhibition of hepatocyte microRNA activity using tough decoys

To perform large-scale miRNA inhibition screens in the repopulating mouse liver, we developed an *in vivo* tough decoy (TuD) expression platform and applied it to the *Fah*^−/−^ mouse model. We generated expression plasmids containing the mouse *Fah* cDNA as well as U6 promoter-driven TuD cassettes (Figure 1B). Each *Fah*-TuD plasmid also encodes the Sleeping Beauty (SB) transposase and transposable elements, facilitating stable integration of the *Fah* and TuD expression cassettes into the hepatocyte genome. Expression of shRNAs from strong RNA Pol III promoters can induce hepatotoxicity *in vivo* by oversaturating the exportin-5 pathway. ^20^ Because RNA Pol III-driven TuD expression might also be toxic to hepatocytes, we designed a second TuD expression cassette to attenuate transcript levels, by removing the termination signal (poly(T); ‘pT’) immediately adjacent to the TuD sequence to allow transcription to proceed to a downstream termination signal. By using two different RNA Pol III expression cassettes, we speculated that at least one would not induce hepatotoxicity. Ultimately, both plasmid systems allowed for effective liver repopulation without triggering excess liver injury, thereby serving as parallel replicate experiments.

To determine if a single TuD can inhibit the activity of up to two different miRNAs in the context of *Fah* transgene expression, we performed dual-luciferase reporter experiments in Hepa 1-6 hepatoma cells *in vitro*. Single- or dual-targeting TuDs directed against members of the miR-194/192/215 family and expressed from either of the two *Fah*-TuD plasmids significantly increased the ratio of *Renilla* luciferase (RLuc) to firefly luciferase (FLuc) activities (Figure 1C). The observed increases in RLuc activity are consistent with the inhibition of miRNA-directed RISC targeting of binding sites within the RLuc 3’UTR. Notably, dual-targeting TuDs displayed miRNA inhibition comparable in magnitude to the associated one-wise TuD constructs, suggesting that TuD levels were not limiting in either of our expression platforms. As expected, TuDs designed to target miR-192 also showed significant de-repression of a miR-215 sensor transcript due to the high similarity between the two mature miRNA sequences. In addition, no effect of TuD expression on luciferase activity was observed in cells expressing a control RLuc without miRNA binding sites. Based on these findings, we reasoned that TuDs can be employed to effectively inhibit endogenous hepatocyte miRNAs in a pairwise manner *in vivo*.

### Generation of pairwise tough decoy libraries for a massively parallel microRNA inhibition screen

To perform a massively parallel screen of miRNA function during liver repopulation, we began by selecting the most relevant miRNAs to be targeted. We utilized our previously-published dataset of miRNAs bound by Argonaute2 (AGO2) in the quiescent mouse liver and following partial hepatectomy^3^ to establish a miRNA-binding site (MBS) pool. For inclusion in our MBS pool, we set a minimum threshold of 0.01% of total AGO2-bound miRNA reads at any of four time points during liver regeneration (quiescent liver and 1, 36, and 48 h after partial hepatectomy). We excluded miRNAs containing a poly(A) stretch of length four or greater, as a corresponding MBS would include a RNA Pol III termination signal. ^21^ All together, we included a total of 171 confidently annotated mature hepatocyte-relevant miRNAs in our library. MBSs targeting three scrambled-sequence miRNAs (scramble-1, −2, −3) and three miRNAs present below two reads per million in the partial hepatectomy dataset (miR-1a-1-5p, miR-670-3p and miR-880-5p) served as negative controls, giving a total pool size of 177 MBSs (Supplementary Table 1). The TuD library was assembled *en masse* via sequential ligation of the MBS pool to a storage plasmid containing TuD stem and loop sequences (Figure 2a and Supplementary Figure 1). This TuD library, containing up to 31,329 possible unique pairwise combinations of 177 unique MBSs, was subcloned into our two *Fah* expression plasmids to test their effect on hepatocyte repopulation (Figure 2A).

**Figure 2.**
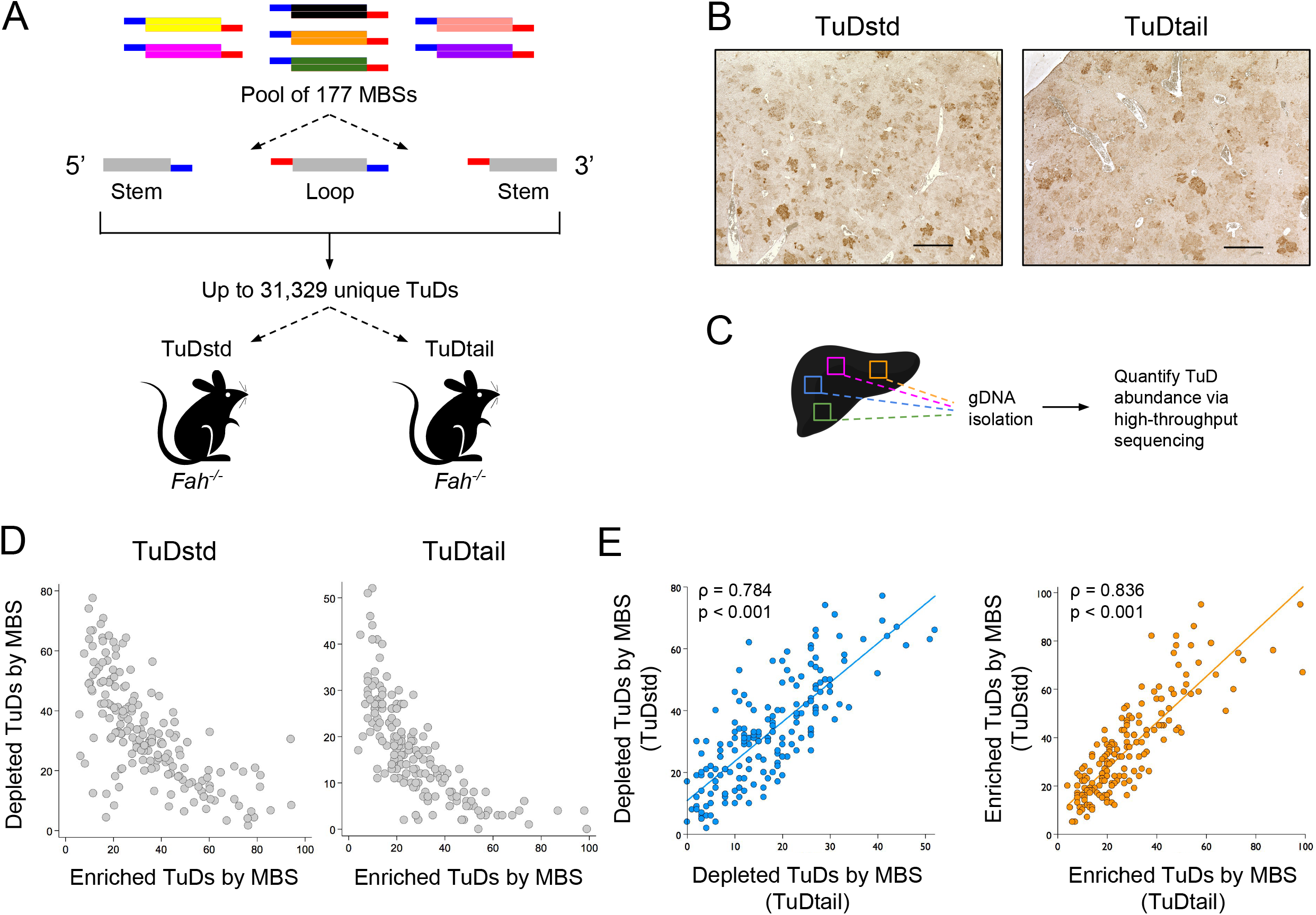
High-content pairwise miRNA inhibition *in vivo*. (A) Schema for pairwise miRNA inhibition screen during liver repopulation. Fah/TuD plasmid libraries were administered to male *Fah*^*−/−*^ mice and livers analyzed four weeks later. (B) Immunohistochemical analysis of *Fah*^*−/−*^ mouse liver tissue. Numerous FAH-positive repopulation nodules are apparent. Scale bar, 500 μm. (C) Schematic of TuD quantification. (D) MBS tallies in enriched and depleted TuDs were negatively correlated in the TuDstd and TuDtail experiments. (E) Depleted (left panel) and enriched (right panel) MBS tallies were significantly correlated between experiments. □, Spearman’s rank correlation coefficient.

### High coverage of tough decoy libraries in the regenerating mouse liver

The TuD plasmid pools, termed ‘TuDstd’ (with pT) or ‘TuDtail’ (without pT) were administered to adult male *Fah*^−/−^ mice via hydrodynamic tail vein injection and nitisinone was withdrawn to induce liver repopulation by hepatocytes incorporating the *Fah*-TuD transposons. Four weeks after plasmid injection, mice were euthanized and liver tissue was collected for analysis. Immunohistochemical analysis confirmed an abundance of FAH-positive repopulation nodules in plasmid-injected *Fah*^−/−^ mice (Figure 2B).

To quantify TuD abundance, we generated sequencing libraries of TuD fragments amplified from repopulated liver genomic DNA (gDNA) and from the input plasmid DNA (Figure 2C). Amplified TuD fragments included both miRNA-binding sites, allowing for TuD identification from a single sequencing read. Our TuDstd and TuDtail plasmid libraries contained 30,817 and 30,846 (98.4% and 98.5%), respectively, of the 31,329 possible pairwise MBS combinations, indicating high cloning efficiency. Following liver repopulation, 30,317 and 30,796 (96.8% and 98.3%) TuDs were detected in liver gDNA samples (Supplementary Table 2). Notably, we observed significant correlations among replicates of the plasmid libraries and liver samples, attesting to the robustness of the assay (Supplementary Figure 2). These data demonstrate that very large combinatorial or genome-scale screens can be performed *in vivo* using the *Fah*^−/−^ mouse model.

### MicroRNAs regulate hepatocyte proliferation *in vivo* following toxic liver injury

Principal component analysis of sequencing reads from each TuD experiment showed a clear separation of liver and plasmid input replicates, suggesting that inhibition of specific miRNAs elicits substantial effects on hepatocyte repopulation (Supplementary Figure 2). We used the differential TuD abundance between plasmid input libraries and hepatocyte gDNA as a measure of miRNA impact on liver repopulation. To identify TuDs with significantly altered abundance, we performed differential expression analysis using DESeq2. ^19^ In TuDstd-injected animals, 5,969 TuDs (19.5%) had an adjusted p-value < 0.05, whereas in TuDtail-injected animals, 3,981 TuDs (13.0%) were significantly altered in abundance after repopulation (Supplementary Figure 3). Many MBSs displayed a strong bias towards either enrichment or depletion; indeed, individual MBS prevalence among the enriched and depleted TuD sets was inversely correlated (Figure 2D).

**Figure 3.**
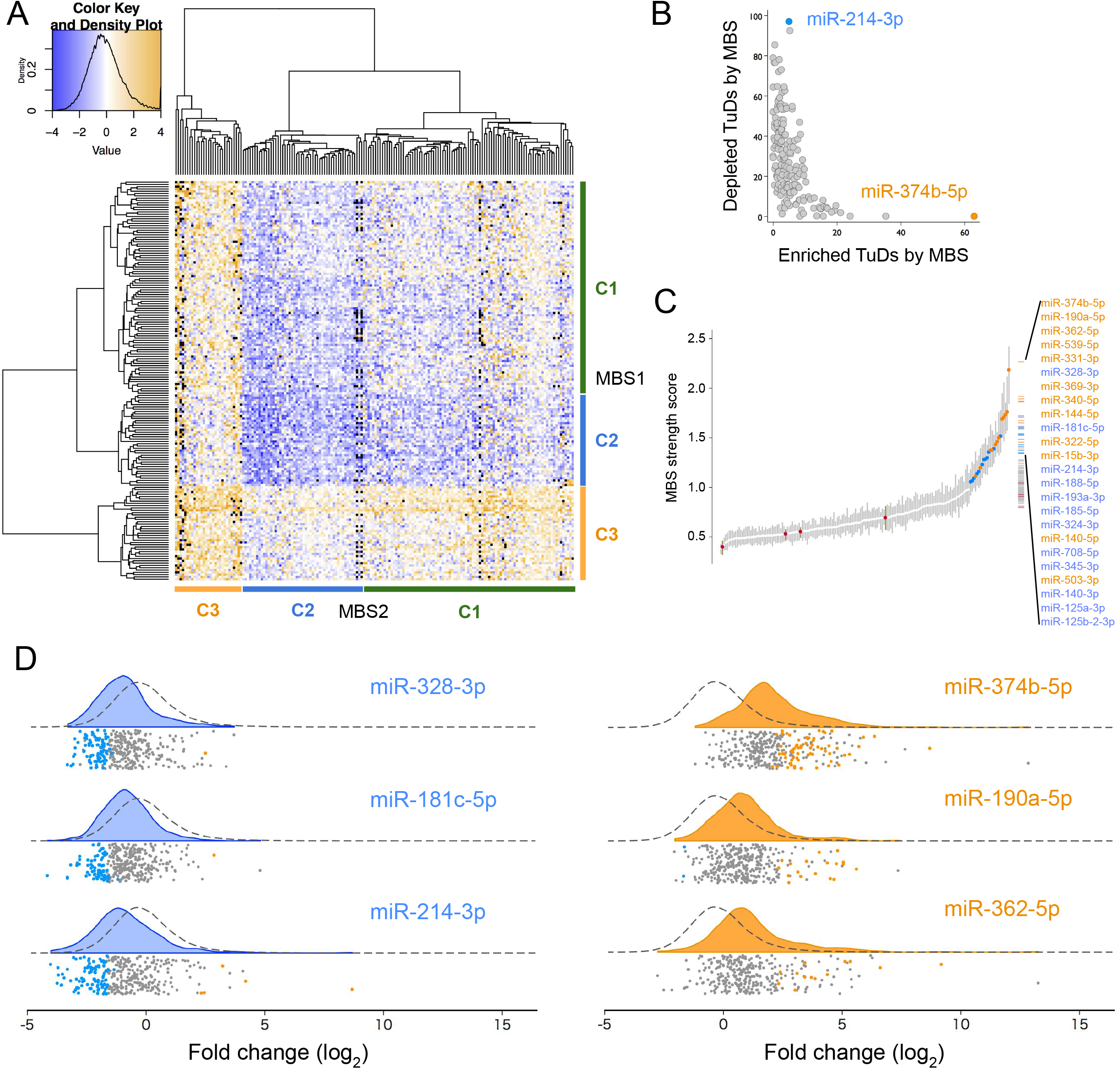
MicroRNAs regulate repopulation following liver injury. (A) Dendrograms and heatmap of log_2_ fold changes of the combined datasets. Colored bars indicate cluster annotations; black, missing values. (B) MBS tallies in enriched and depleted TuDs were negatively correlated in the TuDstd and TuDtail experiments. The most prevalent MBSs in significantly altered TuDs are labeled. (C) Quantile plot of 174 individual MBS strength scores. Control MBS scores are shown in red. Vertical bars indicate 95% confidence intervals. Twenty four MBSs with strength scores >1 are colored according to phenotypes above (orange) or below (blue) zero. (D) Raincloud plots of the log_2_ fold change distributions of TuDs containing the strongest MBSs with a negative phenotype (blue) and or a positive phenotype (orange). Kernel density plots are shaded according to the MBS phenotype. Dashed lines indicate the density plot of all TuDs not containing the specified MBS. Data points below the density plots indicate each TuD detected containing the specified MBS and are colored by the fold change. Depleted (blue), enriched (orange), or unchanged (gray) relative to non-targeting controls. All displayed distributions were significantly different from the distribution of remaining TuDs (two-sample Kolmogorov-Smirnov test p<0.001).

Next, we compared the results of our two replicate experiments. TuD fold change distributions were significantly correlated between libraries (p<0.001) (Supplementary Figure 3). In addition, the log_2_ fold-change distribution interquartile ranges were similar between the two experiments (0.60 and 0.56). The congruent overlap of enriched and depleted TuD sets was significantly greater than expected by chance (Supplementary Figure 3), and the tallies of enriched and depleted TuDs for each MBS were significantly correlated between libraries (Figure 2E). These findings suggested that the presence of the immediate pT termination signal in the TuDtail experiment did not adversely affect TuD activity *in vivo*, and that the findings of our pairwise miRNA inhibition screen were reproducible. To streamline subsequent analyses, we combined the two datasets. After removal of non-biological variation, correlation among all replicates was improved as expected (Supplementary Figure 3).

We identified significantly altered TuDs in the combined data set by comparing the log_2_ fold change values among animal replicates to the population of control TuDs (i.e. TuDs with a control MBS in both MBS positions). After correction for multiple testing, a total of 3,077 TuDs were significantly altered following repopulation, 2,579 (84%) of which were depleted (Supplementary Figure 3). The most prevalent MBSs among depleted and enriched TuDs were directed against miR-214-3p and miR-374b-5p, respectively (Figure 3b), in agreement with our analysis prior to data set combination.

To group MBSs according to their effects on liver repopulation, we performed hierarchical clustering of the TuD log_2_ fold change values (Figure 3A). As each MBS pairing can occur in two orientations (5’-AB-3’ and 5’-BA-3’), our heatmap is MBS order-specific, with rows indicating the 5’ position, and columns representing the 3’ position. In each orientation, three main clusters emerged. We assigned cluster numbers based on the trend towards enrichment or depletion. The cluster 2 MBS sets displayed a strong bias towards depleted TuDs, whereas the cluster 3 MBS sets were biased towards enrichment.

The intersection of the cluster sets across MBS position (Pearson’s □^2^=45.4, p<0.001; Supplementary Figure 4) demonstrated that MBS effects on liver repopulation were largely position-independent. Furthermore, we examined the effect of MBS orientation for all pairwise MBS combinations and observed a significant positive correlation of log_2_ fold change between TuD pairs (5’-AB-3’ and 5’-BA-3’) (Supplementary Figure 4). In addition, each set of significant TuDs was more likely to contain both orientations of an MBS combination than would be expected by chance (Supplementary Figure 4).

**Figure 4.**
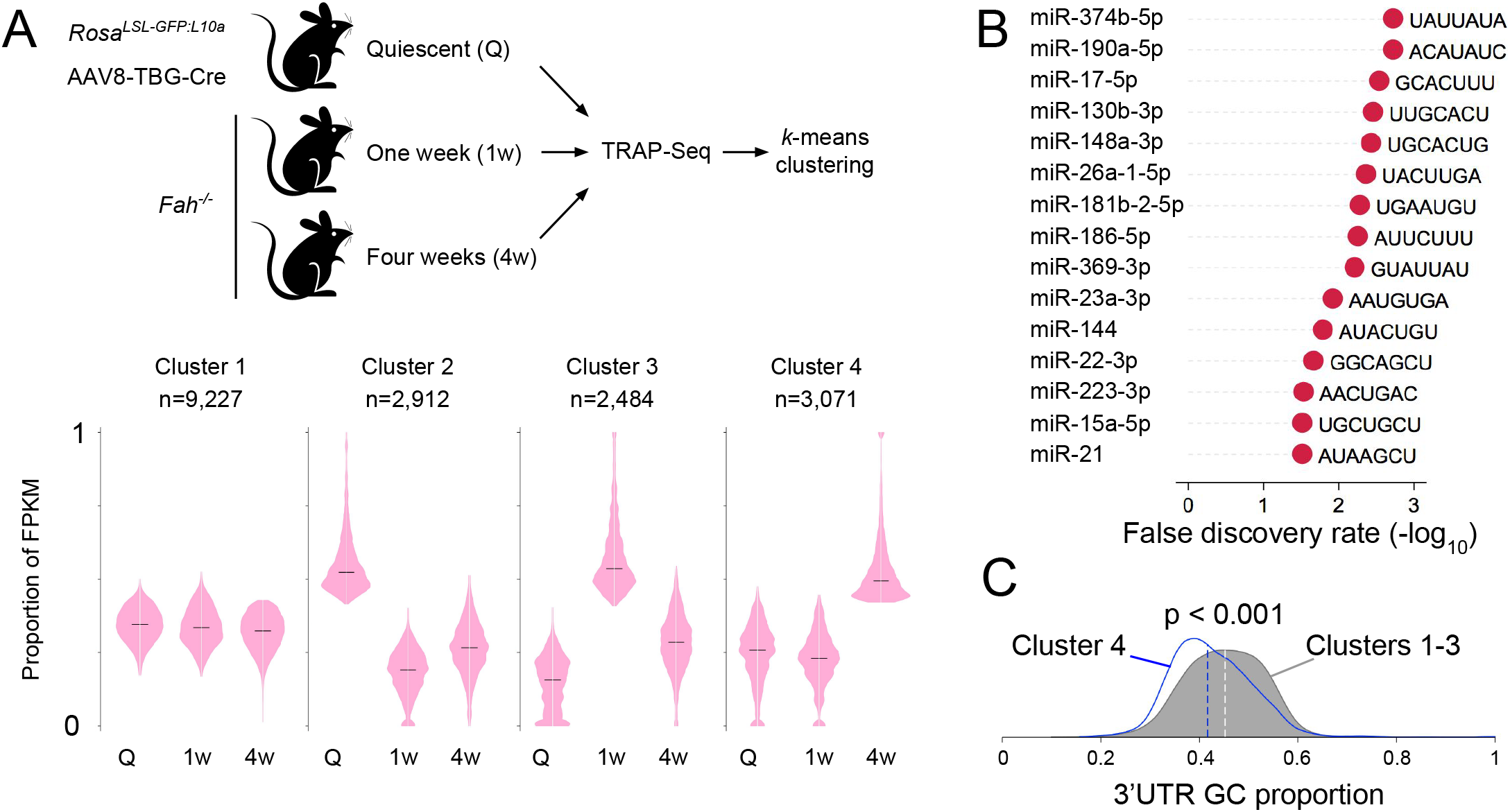
Putative miRNA regulation of the dynamic transcriptome of repopulating hepatocytes. (A) Messenger RNAs from Wang et al. data set were clustered by *k*-means partitioning into four groups. (B) The 3’UTRs of cluster 4 mRNAs were significantly enriched for seed recognition sequences of several miRNAs identified as strong regulators of liver repopulation. (C) Kernel density plots of mRNA and 3’UTR GC content. Dashed vertical lines indicate population medians. Cluster 4 transcripts have significantly reduced GC content as assessed by a two-sample Kolmogorov-Smirnov test.

### Defining individual miRNA binding site effects

Our pairwise screening approach allows for the characterization of individual MBS effects on liver repopulation when functioning alone and when combined with a second non-control MBS. We assigned to each MBS a ‘phenotype’ value that quantifies the effect of inhibiting a single miRNA during liver repopulation. This MBS phenotype was defined as the median log_2_ fold change among the subset of TuDs containing the MBS paired with one of six control MBSs, across all replicates. The lowest phenotype score was observed for the MBS targeting miR-214-3p, which was the most prevalent MBS within depleted TuDs (Supplementary Figure 3). Similarly, the MBS targeting miR-374b-5p, the most prevalent MBS among enriched TuDs, was assigned the second highest phenotype score (Supplementary Figure 3).

Next, we developed a metric to capture the ability of an MBS to affect liver repopulation in the presence of a second MBS. For each TuD, we assigned two MBS ‘strength scores’ based on the constituent MBS phenotypes and the observed TuD log_2_ fold-change. Each strength score is the ratio of the distance between MBS phenotypes and the distance between the MBS phenotype and the observed log_2_ fold-change (Supplementary Figure 3). For each MBS, an overall strength score was calculated as the median strength score for the MBS across all TuDs and replicates. The largest overall strength score was observed for miR-374b-5p (Figure 3C). Of note, this approach allowed for comparison of the magnitude of the impact of each MBS on liver repopulation, regardless of whether MBS phenotypes enhance or inhibit regeneration (Supplementary Figure 3). We then examined the log_2_ fold-change distributions of the strongest MBSs with positive (enriched) or negative (depleted) phenotypes and found these distributions to be significantly different than the population of TuDs not containing the MBS of interest (Kolmogorov-Smirnov p-value < 0.001) (Figure 3D).

We compared our strength metric with the Bradley-Terry model of pairwise comparisons. ^22^ For each pairwise TuD (i.e. having two different MBSs) we assigned a ‘win’ or ‘loss’ to each constituent MBS based on MBS phenotypes and the observed TuD log_2_ fold-change (Supplementary Figure 5). Win and loss tallies for each combination across all replicates were then used to derive model coefficients for each MBS. In line with our strength score, the Bradley-Terry model classified the miR-374b-5p MBS as the most potent regulator of repopulation, and Bradley-Terry model coefficients were significantly correlated with our MBS strength metric (Supplementary Figure 5). Altogether, these analyses classified a subset of MBSs as modifiers of hepatocyte repopulation following toxic liver injury, implicating specific miRNAs in post-transcriptional regulation of liver repopulation.

**Figure 5.**
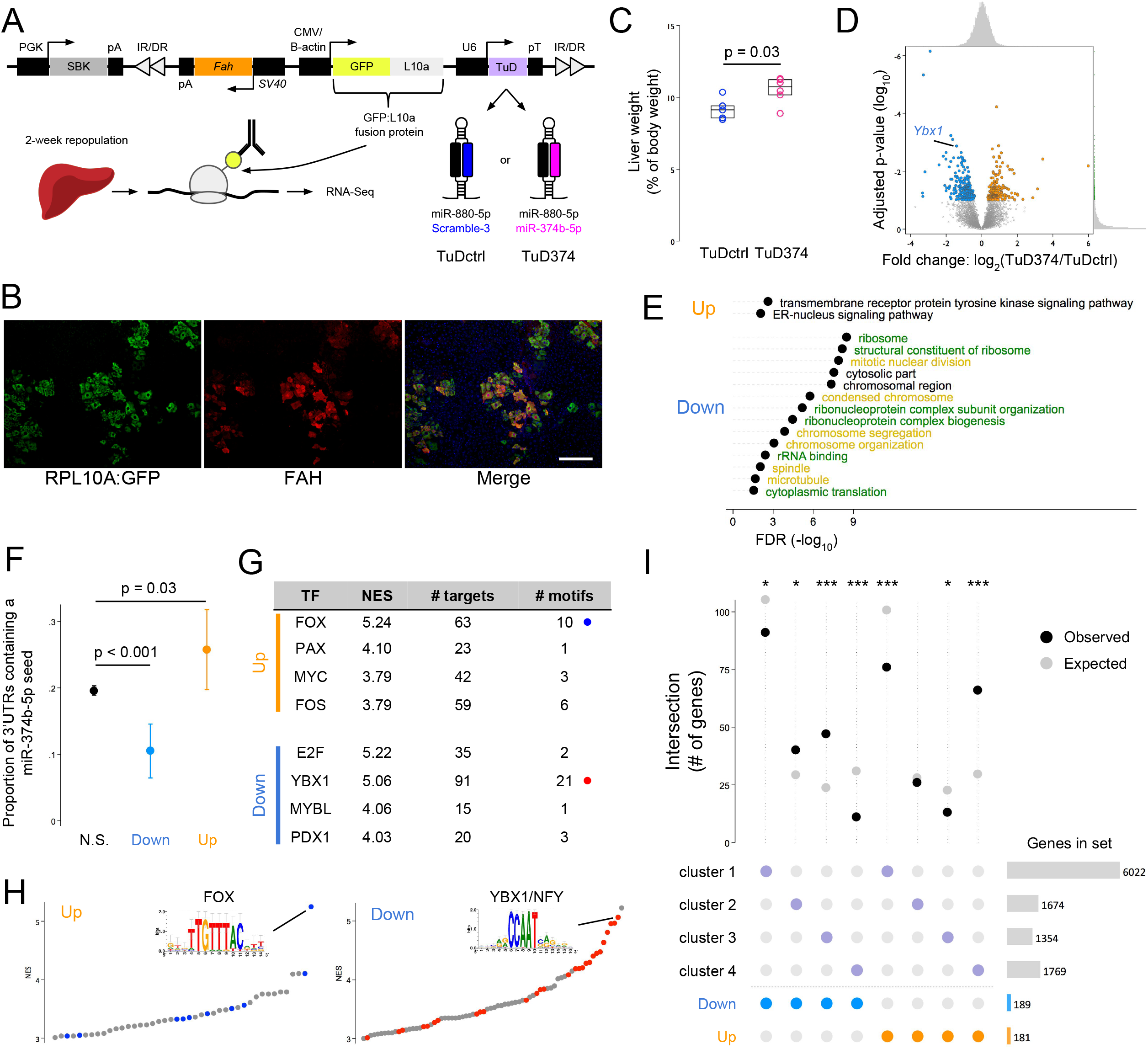
Inhibition of miR-374b accelerates liver repopulation. (A) Profiling the miR-374b-5p target transcriptome using TRAP-Seq. *Fah* plasmids co-expressing an RPL10A:GFP fusion protein and a TuD targeting miR-374b-5p or control miRNAs were delivered to *Fah*^*−/−*^ mice. Translating ribosomes were isolated after two weeks of repopulation and ribosome-bound mRNAs quantified by next generation sequencing. (B) Immunofluorescent staining of liver sections confirming co-expression of the RPL10A:GFP fusion protein (green) and FAH (red). DAPI, blue; scale bar, 200 μm. (C) Liver weight as a percentage of total body weight. Box overlays indicate the sample median and interquartile range. (D) Differential ribosomal occupancy between groups as determined by DESeq2. (E) Gene ontology overrepresentation analysis of the differentially expressed transcript sets. Terms associated with the ribosome (green) or cell replication (yellow) are colored. (F) Proportion of 3’UTRs containing a miR-374b-5p seed recognition site. Data are presented as mean ± 95% CI. (G) Motif analysis of genes up- or down-regulated by TuD374 within +/-20 kilobase of the TSS. Transcription factors (TF) associated with the top four motif clusters by normalized expression score (NES) within each gene set are shown. (H) Quantile plots of enriched motif NES. Colored data points indicate enriched motifs for the indicated motif cluster. Sequence logos of the top motifs are shown as insets. (I) Upset plot of the intersect between differentially expressed genes of the TuD TRAP-Seq experiment and the gene clusters of Figure 4B. TuD374 livers were enriched for transcripts elevated at 4-weeks (cluster 4) and depleted for transcripts elevated at week 1 (cluster 3). *p<0.05, ***p<0.001.

### Hepatocyte miRNAs regulate diverse pathways

To examine transcripts induced in FAH-positive repopulating hepatocytes for putative miRNA binding sites, we utilized our previously published TRAP-Seq data set generated using the *Fah*^*−/−*^ mouse model. ^2^ Transcripts in repopulating hepatocytes were assigned to one of four clusters using *k*-means clustering based on temporal changes in expression level (Figure 4A). The 3’UTRs of transcripts of cluster four (n=3,071), which displayed elevated expression at four weeks after the initiation of liver injury, were significantly enriched for the seed recognition sequences of several miRNAs identified in our screen as strong regulators of repopulation. Remarkably, the two most significantly enriched seeds were those of miR-374b-5p and miR-190a-5p, the targets of the two strongest MBSs (Figure 4B). As a group, the miRNAs with enriched seed abundance in cluster four showed a high proportion of A or U nucleotides (65.7%) within the seed sequence. Interestingly, the mRNAs of cluster four had significantly higher A/U-content compared to transcripts of the remaining three clusters (Supplementary Figure 6); this observation was also true when we restricted the analysis to 3’ UTRs (Figure 4C). Repopulating hepatocytes may therefore upregulate a subset of miRNAs to coincide with this shift in the protein-coding transcriptome.

**Figure 6.**
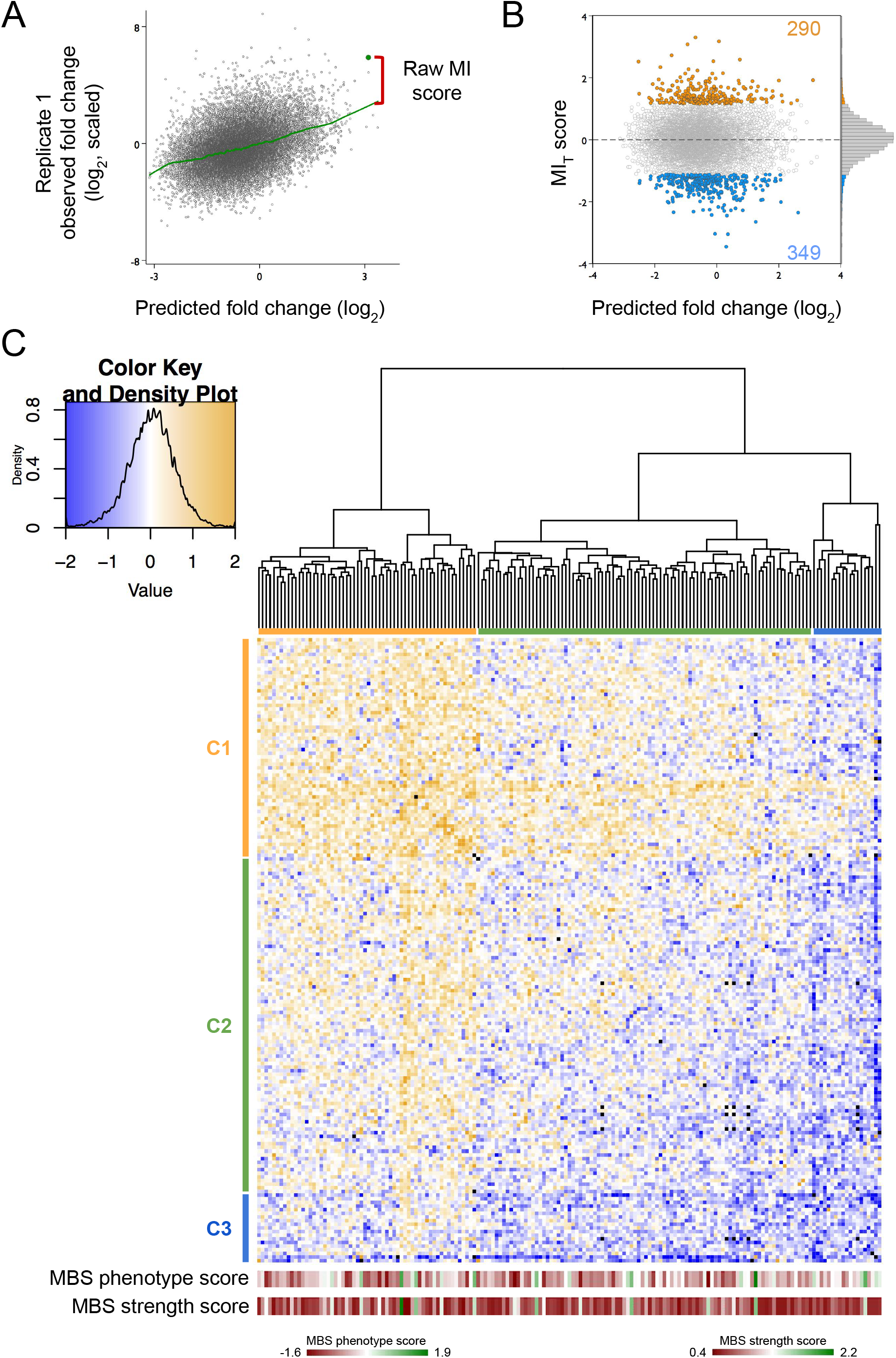
Epistatic miRNA interactions uncovered by pairwise TuD screens. (A) Raw miRNA interaction (MI) scores were calculated as the residuals from a LOWESS fit line of the observed versus predicted TuD fold change plot. (B) Scatterplot of modified t-scores of raw MI data (MI_T_ score) and predicted log_2_ fold changes. MI_T_ scores at least two standard deviations above or below the population mean were considered significant interactions. (C) Hierarchical clusters and heatmap of MI_T_ scores for all TuDs. Individual MBS phenotypes and strength scores are shown in rows below the MI_T_ map.

### MicroRNA-374b inhibition in hepatocytes accelerates liver repopulation

Our previously published report of AGO2-bound miRNAs found that liver miR-374b-5p levels increase significantly following partial hepatectomy (Supplementary Figure 6). ^3^ We also noted that the promoter of the miR-374b host gene, *Ftx*, exhibited increased chromatin accessibility following the induction of liver repopulation in *Fah*^*−/−*^ mice (Supplementary Figure 6). Taken together, these results suggested that the miR-374b-5p target transcriptome in hepatocytes includes genes important for liver repopulation following hepatotoxic injury.

To test this hypothesis, we developed Fah-TuD co-expression plasmids that facilitate TRAP-Seq analysis in the presence of specific miRNA inhibition (Figure 5A). Each TRAP plasmid encodes a fusion protein consisting of the ribosomal protein RPL10A and enhanced GFP, along with a TuD targeting miR-374b-5p (miR-880-5p paired with miR-374b-5p, ‘TuD374’) or control miRNAs (miR-880-5p paired with scramble-3, ‘TuDctrl’). TRAP plasmids were administered to *Fah*^−/−^ mice and liver repopulation was allowed to proceed for two weeks prior to tissue collection. Immunofluorescent imaging confirmed the co-expression of FAH and the RPL10A:GFP fusion protein in repopulating hepatocytes (Figure 5B). The liver to body weight ratio at the time of collection was significantly higher in animals treated with TuD374 compared to TuDctrl, despite similar overall body weights between groups (Figure 5C and Supplementary Figure 7), indicating that repopulation had occurred more robustly in the TuD374 group.

**Figure 7.**
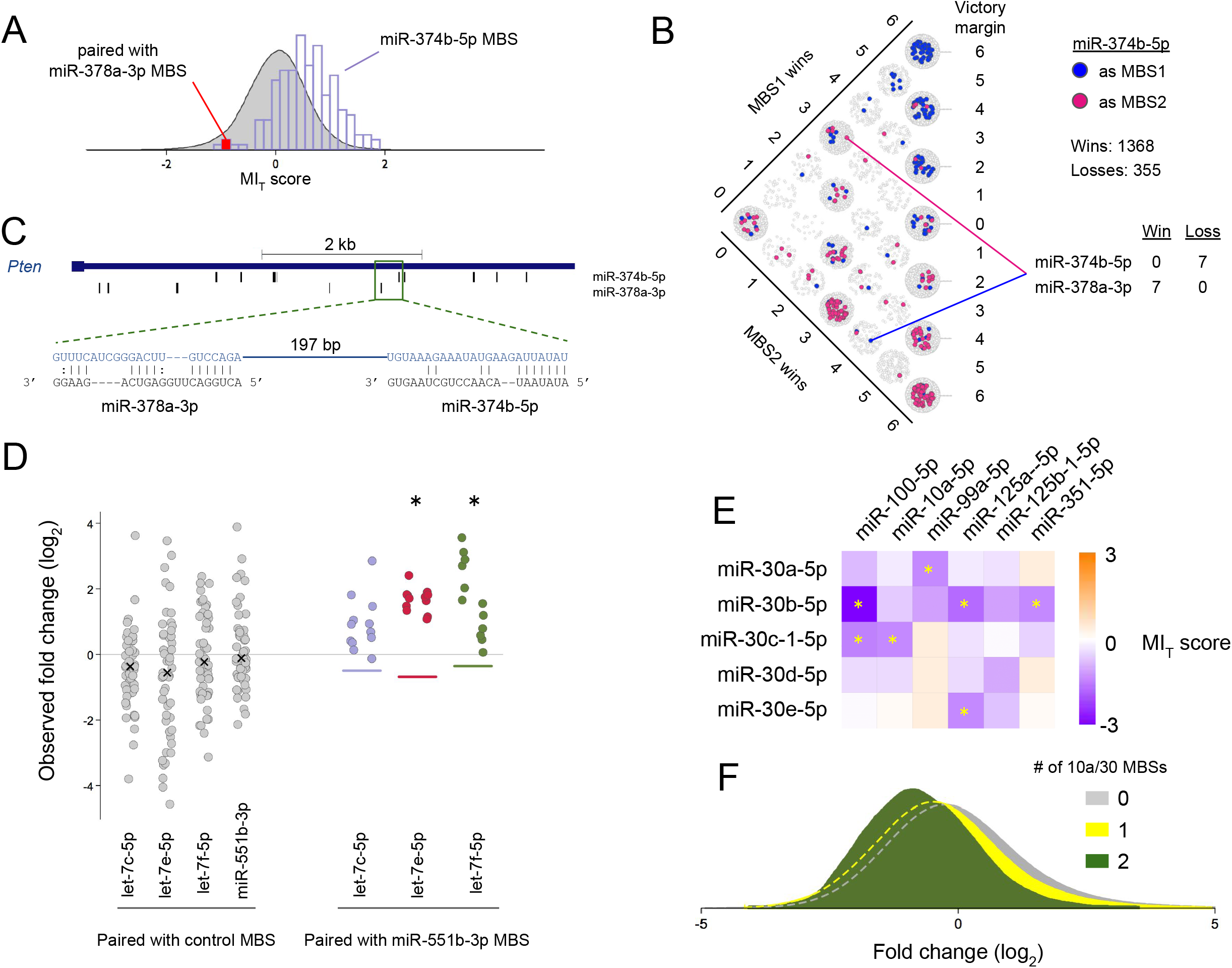
Examples of epistatic interactions between miRNAs. (A) The distribution of MI_T_ scores of TuDs containing a miR-374b-5p MBS (lavender histogram) is shifted right compared to the full TuD population (gray distribution). The MI_T_ score of the miR-378-3p pairing is among the lowest for TuDs containing a miR-374b-5p MBS. (B) Mancala plot of TuDs containing a miR-374b-5p MBS. Despite a high winning percentage across all combinations, miR-374b-5p collected zero wins when paired miR-378-3p. (C) Schematic of *Pten* transcript 3’UTR and putative miRNA binding sites for miR-374b-5p and miR-378-3p. (D) The observed fold changes for TuDs containing a control MBS paired with an MBS targeting members of the let-7 family or miR-551b-5p. Each data point represents a fold-change from a single mouse replicate. The expected fold-changes for pairwise combinations are indicated by colored lines. Data points for each pairwise TuD (colored) are separated horizontally by orientation (AB, left; BA, right). Asterisks indicate a significant interaction by MI_T_ score. (E) Heatmap of MI_T_ scores of pairwise TuDs targeting the miR-10 and miR-30 families. Asterisks indicate significant interaction by MI_T_ scores. (F) Kernel density plots of log_2_ fold change values of TuDs containing the indicated number of miR-10 and miR-30 family MBSs.

To assess changes in the hepatocyte transcriptome induced by miR-374b-5p inhibition, we isolated ribosome-bound mRNA from TuD374 and TuDctrl livers via affinity purification and performed RNA-Seq. Differential expression analysis identified 421 genes as significantly altered (FDR < 10%; Figure 5D). Among differentially expressed genes, we noted a striking decrease in transcripts of ribosomal subunit proteins (Supplementary Figure 7) and cell cycle regulators in TuD374 animals. Indeed, gene ontology (GO) analysis found that the set of genes significantly reduced by miR-374b-5p inhibition was enriched for many GO terms associated with cell proliferation and the cytosolic ribosome (Figure 5E). Strikingly, each of these altered ribosomal genes showed a decrease in expression over time in our previous TRAP-Seq time course (Supplementary Figure 7). In addition, several genes highly expressed in fetal liver cells, including *Igf2* and *Afp*, were significantly reduced in TuD374 livers (Supplementary Figure 7). These results suggest that hepatocytes expressing TuD374, while having an increased ability to repopulate, do not express progenitor-like markers.

Transcripts increased in TuD374-expressing hepatocytes were significantly more likely to contain a miR-374b-5p seed sequence in the 3’ UTR compared to unaffected mRNAs (Figure 5F). Several transcripts, including desmocollin 2 (*Dsc2*), were previously identified as miR-374b-5p targets. ^3^ However, the majority of the elevated mRNAs did not contain a miR-374b-5p seed, and we sought to identify indirect mechanisms to account for the observed expression changes. We used iRegulon^23^ to identify transcription factor binding motifs enriched within a 20 kb window centered on the transcriptional start sites of genes in the altered sets. The motif cluster with the highest Normalized Enrichment Score (NES) in the TuD374 upregulated gene set contained forkhead box (FOX) transcription factor recognition sequences (Figure 5G and H). Elevated FOX factor activity suggests that TuD374-expressing hepatocytes had already regained normal metabolic functions following two weeks of accelerated repopulation. Among the most downregulated transcripts in TuD374 hepatocytes was the mRNA encoding Y box protein 1 (YBX1), a transcription factor and RNA-binding protein that interacts with the highly-conserved Y/CCAAT box. This motif, also bound by the nuclear transcription factor Y (NFY) complex, was the most enriched binding motif of the downregulated gene set (Figure 5G and H).

The observed increase in liver size combined with a less proliferative expression profile suggested that TuD374 had accelerated liver repopulation in the two weeks prior to tissue harvest. To further explore this possibility, we compared the sets of significantly altered transcripts to the *k*-means clusters (Figure 4A) we derived from the repopulation time-course TRAP-Seq data. We found a 2.3-fold overrepresentation of genes elevated in TuD374 livers within TRAP-Seq cluster four, which consists of genes increased after four weeks of repopulation, and a 1.7-fold underrepresentation within cluster three, which showed elevation at the one-week time point (Figure 5I). Conversely, genes reduced in TuD374 livers were underrepresented 2.8-fold in cluster four and 2-fold overrepresented in cluster three. These results support the notion that TuD374 hepatocytes had more rapidly executed their repopulation program.

### Epistatic effects of pairwise microRNA inhibition during liver repopulation

Our screening platform was also designed to identify pairwise miRNA interactions, as the binding of multiple RISC to a transcript can elicit cooperative repression to an extent greater than the sum of the individual complexes. ^24^ To uncover epistatic effects, we derived a miRNA-interaction (MI) score for each TuD based on a previously described model of combinatorial effects. ^25^ The raw MI score of a TuD was defined as its residual from a smoothed regression of observed and predicted phenotypes, where the predicted TuD phenotype is equal to the sum of the constituent MBS phenotypes (log_2_ scale) (Figure 6A). Raw MI scores were converted to modified t-value scores (MI_T_ scores) ^26^ using replicate raw MI scores of each MBS pairing in either orientation (i.e. 5’-AB-3’ and 5’-BA-3’). We defined TuDs with MI_T_ scores at minimum two standard deviations from the population mean as displaying significant miRNA interactions resulting in 639 MBS pairings (4.5%) with significant MI_T_ scores (Figure 6B). We next derived a miRNA interaction map by hierarchical clustering of the MI_T_ scores (Figure 6C). As with our fold change map, three main clusters emerged. Overlay of MBS phenotypes or strength scores suggested that clustering on the MI_T_ map was not driven by individual miRNA effects.

Genetic interaction maps, such as our MI_T_ map, are useful in generating hypotheses. ^25,27^ For example, the majority of MI_T_ scores of the miR-374b-5p MBS were positive, but several pairings produced TuD fold change values well below the expected (Figure 7A). The miR-378a-3p MBS pairing had one of the lowest MI_T_ scores for TuDs containing the miR-374b-5p MBS. Win/loss tallies from our Bradley-Terry analysis showed this MBS consistently antagonized the miR-374b-5p MBS (Figure 7B). We hypothesized that miR-378a-3p functions in concert with miR-374b-5p to co-repress a potent negative regulator of hepatocyte repopulation, and that binding of either miR-374b-5p or miR-378a-3p is sufficient to inhibit such a target. Thus, as long as the miR-374b-5p TuD is present in a hepatocyte clone paired with a non-relevant TuD, its growth promoting effect on many other targets is dominant, and the continued repression of the potent and dominant negative regulator is still taken place by binding of miR-378a-3p. A search of putative 3’UTR target sites identified the tumor suppressor *Pten* as containing multiple binding sites for miR-374b-5p and miR-378a-3p (Figure 7C). Thus, removing miR-374b-5p repression of this tumor suppressor alone is insufficient to substantially inhibit repopulation triggered by miR-374b-5p inhibition. We postulate that only when both miR-374b-5p and miR-378a-3p are inhibited does *Pten* mRNA become stabilized and exert a dominant inhibitory effect on liver repopulation.

The highest MI_T_ score was observed for the TuD containing let-7e-5p and miR-551b-3p MBSs. Both of these MBSs had phenotypes below zero, suggesting that let-7e-5p or miR-551b-3p inhibition alone does not enhance liver repopulation. However, co-repression of these miRNAs resulted in consistent enrichment following repopulation (Figure 7D). Strikingly, all TuDs pairing a miR-551b-3p MBS with an MBS targeting let-7c-5p, let-7e-5p, or let-7f-5p had fold-change values greater than expected (Figure 7D). These MBSs did not tightly cluster on our fold-change map, suggesting independent functions due to largely disjoint target mRNA sets. One intriguing candidate for co-regulation by these miRNAs is peroxisomal biogenesis factor 5 (*Pex5*), a sensor of oxidative stress. AGO2 footprints containing sites for miR-551b-3p and let-7e-5p were previously observed on *Pex5* transcripts in regenerating liver cells.^3^

We noted that seven of 30 TuDs (23%) targeting both the miR-30 family and the miR-10 family had significant MI_T_ scores below zero (Figure 7E). The population of TuDs containing two MBSs targeting the miR-30 and/or miR-10 family had significantly lower fold-change values compared to TuDs containing one such MBS or none (Figure 7F). We identified the RAR-related orphan receptor alpha (*Rora*) transcript as a putative target that may account for the observed miRNA interaction. AGO2 footprints on the *Rora* transcript that overlap with miR-30 and miR-10 family recognition sequences. ^3^ These examples illustrate experimental hypotheses generated using our MI_T_ map and the rich resource provided by our *in vivo* pairwise miRNA inhibition screens.

## DISCUSSION

Although genome-scale screens of all possible pairwise combinations of protein-coding transcripts in mammalian cells are currently unfeasible, massive pairwise genetic screens employing shRNA, siRNA, miRNA, or CRISPR expression systems have now been conducted *in vitro*. ^25,27–31^ Given that a mammalian cell type expresses just a subset of the ~1,000 miRNAs encoded in its genome, screens encompassing all pairwise combinations of miRNAs active in a given cell type are now practical. Here we demonstrate a novel approach to screen miRNA function in a pairwise and high-throughput manner *in vivo*.

The continued delivery of miRNA modulators to the liver in an efficacious manner over a repopulation time course is cost-prohibitive. Our sequential ligation approach makes the derivation of large TuD pools cost-effective compared to the synthesis of full-length inhibitors, as the stem and loop sequences shared by all TuDs are incorporated within the plasmid backbone prior to inhibitor assembly. By tailoring our TuD plasmids for use in *Fah*^*−/−*^ mouse model, we have screened inhibitors *in vivo* at a much larger scale than would otherwise be possible. The high proportion of TuDs detected in livers following repopulation confirms that libraries of this complexity can be reliably screened in this model. This is possible because a mouse liver contains ~10^8^ hepatocytes, of which 10^5^-10^6^ are stably transduced by hydrodynamic tail vein injection of transposase plasmids. Due to the strong selective pressure, competitive repopulation by 10^5^-10^6^ hepatocyte clones can be assessed in a single animal. Our system reliably characterized the effects of individual mature miRNAs. The TuD design ensured that a given MBS was present in numerous, uniquely identifiable inhibitors in each library, with up to 353 measurement for each MBS, enabling accurate quantification of effects on repopulation.

The detection of miRNA regulation is often confounded by mild to moderate effect sizes upon individual transcripts. Redundant targeting by multiple miRNAs further complicates the elucidation of their function. High-order systematic miRNA screens afford the opportunity to discover interactions that would otherwise go undetected. ^30^ The results of our pairwise inhibition screens enable the informed design of higher-order miRNA investigations, for which the mouse liver would otherwise be insufficient due to scalability. Exploration of the interaction map generated here may also inform novel combinatorial therapeutic strategies to address aberrant liver cell proliferation. The miRNA inhibition system we developed can be easily adapted to study other conditions for which the *Fah*^*−/−*^ model has been utilized, including hepatocellular carcinoma. ^16^ To our knowledge, this study is the first to systematically test the functions of a mammalian tissue’s miRNAome.

## Notes

**Grant support:** This work was supported by the following awards from the NIH: R01 DK102667 (K.H.K.), K01 DK102868 (A.M.Z.), K08 DK106478 (K.J.W.), and F31 DK113666 (A.W.W.). We thank the University of Pennsylvania Diabetes Research Center for the use of the Functional Genomics Core (P30 DK19525) and the Center for Molecular Studies in Digestive and Liver Diseases (P30 DK050306) for the use of the Molecular Pathology and Imaging Core. We also thank Long Gao for bioinformatics support.

