## Supplementary figures and images for "A high-content *in vivo* screen to identify microRNA epistasis in the repopulating mouse liver"

### Supplementary Figure 1

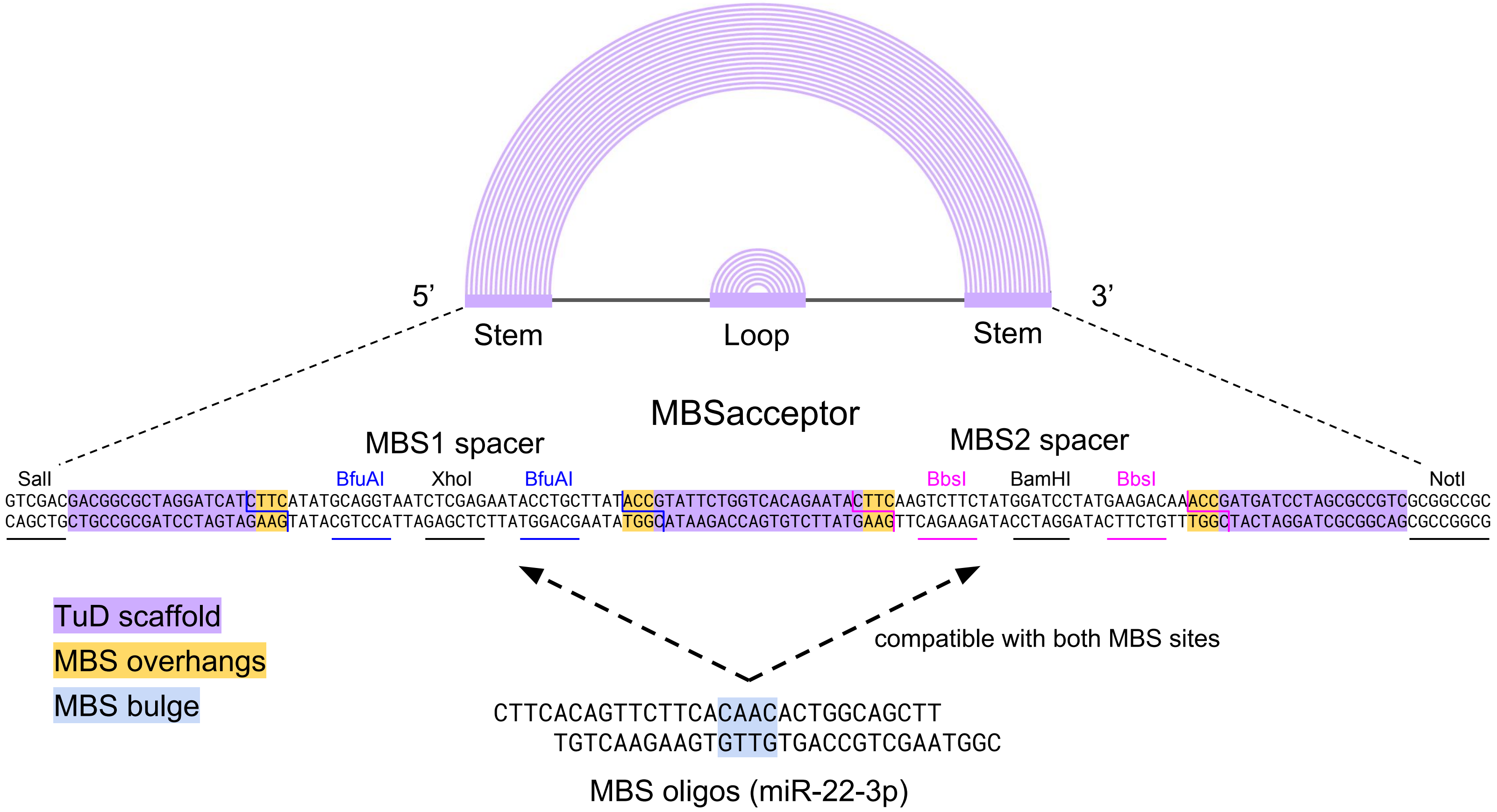


Supplementary Figure 1

### Supplementary Figure 2

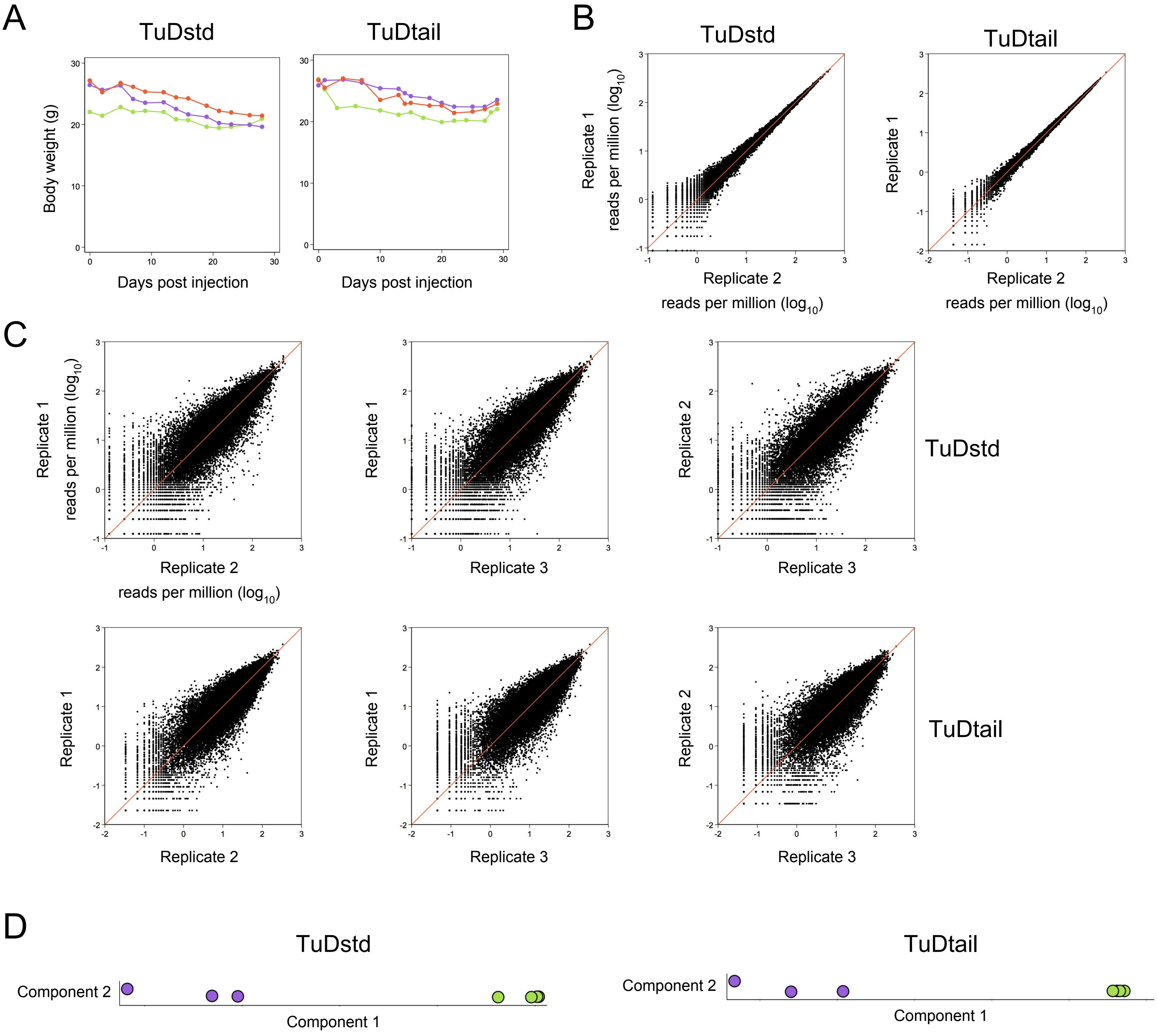


Supplementary Figure 2

### Supplementary Figure 3

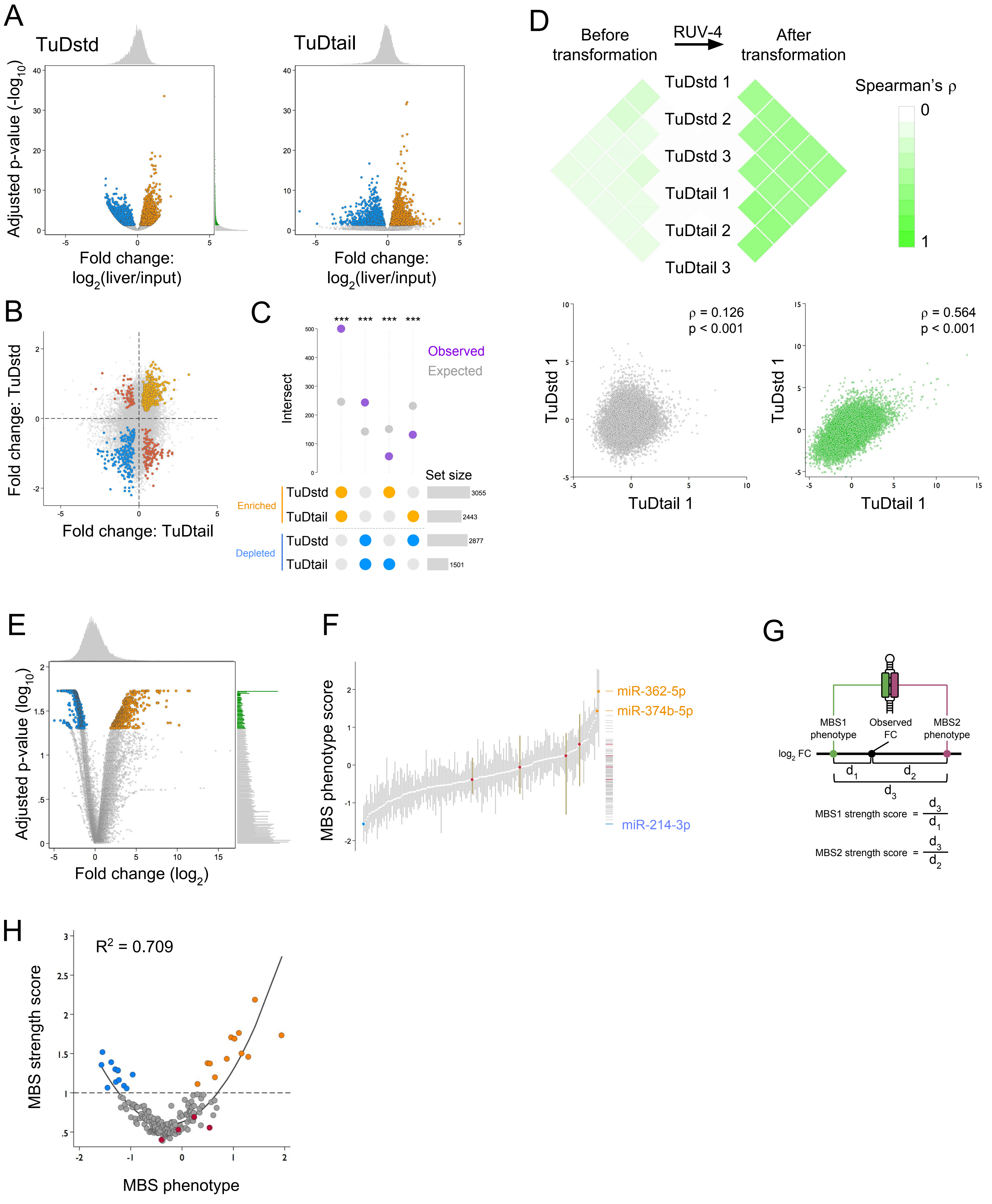


#### Supplementary Figure 3

### Supplementary Figure 4

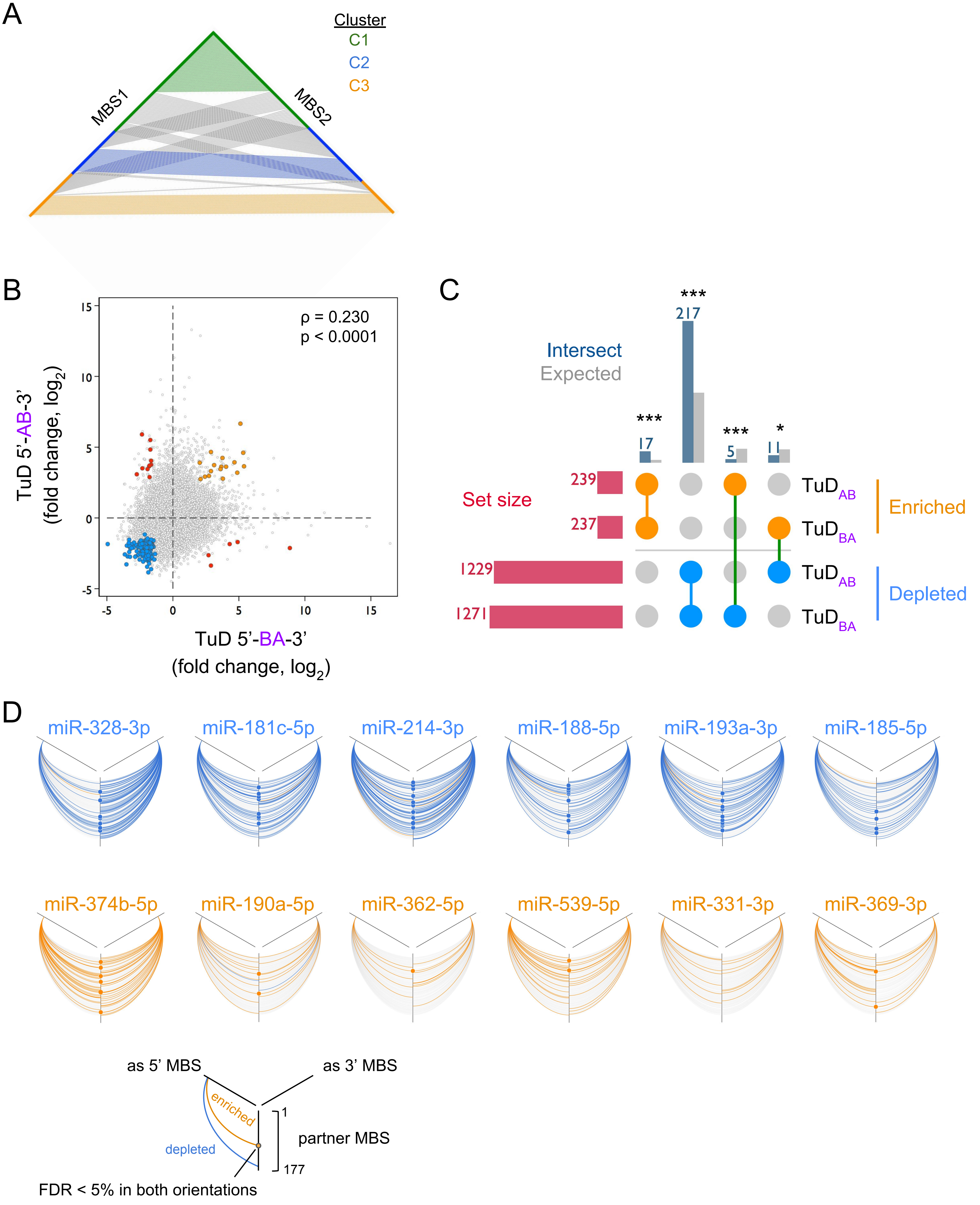


Supplementary Figure 4

### Supplementary Figure 5

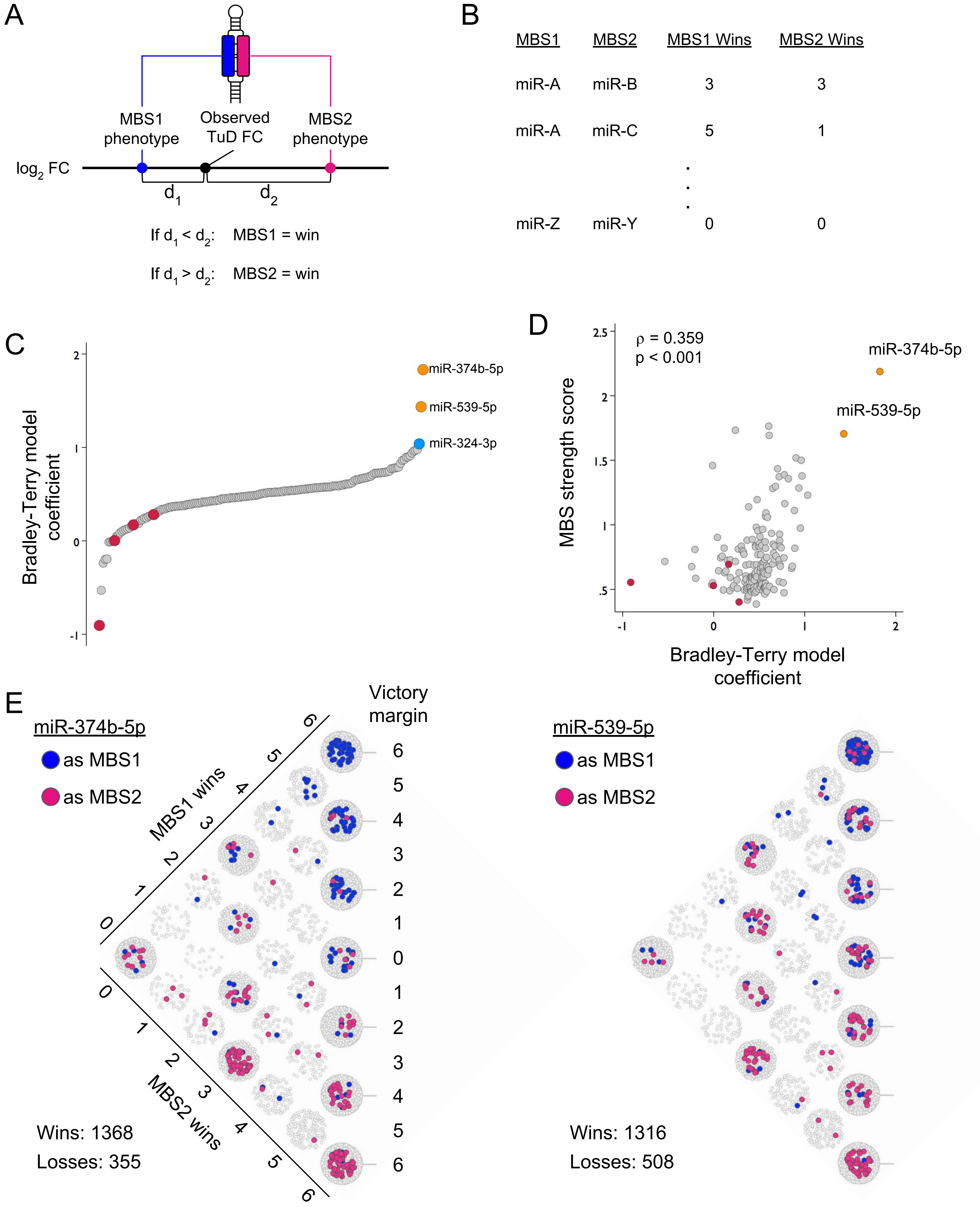


Supplementary Figure 5

### Supplementary Figure 6

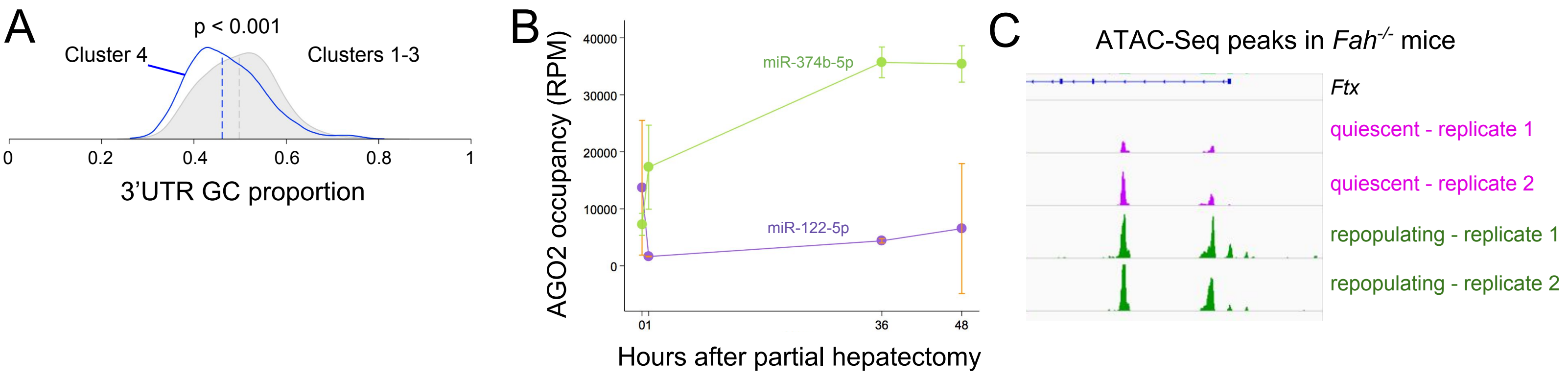


Supplementary Figure 6
